# Variants at *RNF212* and *RNF212B* are associated with recombination rate variation in Soay sheep (*Ovis aries*)

**DOI:** 10.1101/2020.07.26.217802

**Authors:** Susan E. Johnston, Martin A. Stoffel, Josephine M. Pemberton

## Abstract

Meiotic recombination is a ubiquitous feature of sexual reproduction, ensuring proper disjunction of homologous chromosomes, and creating new combinations of alleles upon which selection can act. By identifying the genetic drivers of recombination rate variation, we can begin to understand its evolution. Here, we revisit an analysis investigating the genetic architecture of gamete autosomal crossover counts (ACC) in a wild population of Soay sheep (*Ovis aries*) using a much larger dataset (increasing from 3,300 to 7,235 gametes and from ∼39,000 to ∼415,000 SNPs for genome-wide association analysis). Animal models fitting genomic relatedness confirmed that ACC was heritable in both females (*h*^2^ = 0.18) and males (*h*^2^ = 0.12). Genome-wide association studies identified two regions associated with ACC variation. A region on chromosome 6 containing *RNF212* explained 46% of heritable variation in female ACC, but was not associated with male ACC, confirming the previous finding. A region on chromosome 7 containing *RNF212B* explained 20-25% of variation in ACC in both males and females. Both *RNF212* and *RNF212B* have been repeatedly associated with recombination rate in other mammal species. These findings confirm that moderate to large effect loci can underpin ACC variation in wild mammals, and provide a foundation for further studies on the evolution of recombination rates.

## Introduction

Meiotic recombination is ubiquitous in sexually-reproducing organisms and is an important driver of genomic diversity in eukaryotes (Barton and Charlesworth, 1998; Otto and Lenormand, 2002). The process of crossing-over ensures the proper disjunction of homologous chromosomes during meiosis (Hassold and Hunt, 2001), with most species having a minimum requirement of at least one obligate crossover per chromosome pair (Otto and Payseur, 2019). It also has the advantage of generating novel haplotypes and preventing the accumulation of harmful mutations in the genome, allowing populations to respond to selection faster and to a greater degree (Muller, 1964; Hill and Robertson, 1966; Battagin *et al*., 2016). However, this can come at the cost of increased mutations at crossover sites (Halldorsson *et al*., 2019) and the break up of favourable haplotypes (Barton and Charlesworth, 1998). Variation in recombination rate is likely to arise across taxa with variation in genome size and chromosome number (Stapley *et al*., 2017); in addition, the evolutionary costs and benefits of recombination may vary depending on the selective context, again leading to the expectation of recombination rate variation within and between populations (Otto and Barton, 2001; Otto and Lenormand, 2002). Indeed, cytogenetic and linkage mapping studies have shown that recombination rates can vary by orders of magnitude within and between chromosomes (Myers *et al*., 2005), individuals (Kong *et al*., 2004), populations (Samuk *et al*., 2020), sexes (Lenormand and Dutheil, 2005) and species (Stapley *et al*., 2017).

In mammals, genomic studies in different species (i.e. humans, cattle, sheep, pigs and deer) have shown that recombination rate is often heritable, with a significant proportion of the additive genetic variation underpinned by loci of moderate to large effects (Sandor *et al*., 2012; Kong *et al*., 2014; Ma *et al*., 2015; Kadri *et al*., 2016; Petit *et al*., 2017; Johnston *et al*., 2018; Johnsson *et al*., 2020). Genome-wide association studies (GWAS) have shown repeated associations with variants at ring finger protein 212 *RNF212* and its paralogue *RNF212B*, whose associated proteins are essential for the crossover designation process during meiosis (Reynolds *et al*., 2013), as well as other loci with established roles in meiotic processes (e.g. *HEI10, REC8* and *TOP2*; Sandor *et al*. 2012; Kong *et al*. 2014; Ma *et al*. 2015; Kadri *et al*. 2016; Petit *et al*. 2017; Johnston *et al*. 2018; Johnsson *et al*. 2020). This heritable, oligogenic architecture implies that recombination rate has the potential to evolve rapidly to selection if variation in recombination rate is associated with individual fitness (Otto and Barton, 2001; Stapley *et al*., 2017). Therefore, detailed examination of the genetic architecture of recombination rate is a critical step in understanding how rates are evolving over different evolutionary timescales.

The Soay sheep (*Ovis aries*) are a Neolithic breed of domestic sheep that have been living wild on the St Kilda archipelago since the Bronze age. These sheep are a model system for studies of ecology and evolution, and have been intensively studied since 1985 (Clutton-Brock *et al*., 2004). A previous study integrated pedigree information with genotypic data from ∼39,000 SNPs to characterise autosomal crossover counts (ACCs) in ∼3,300 gametes transmitted from parents to their offspring (Johnston *et al*., 2016). A GWAS showed that a single region on chromosome 6, corresponding to the locus *RNF212*, explained around 47% of heritable variation in female ACC only. A broader-scale “regional heritability” analysis identified a tentative association in both sexes combined at a 1.09 Mb region on chromosome 7, containing the candidate loci *REC8* and *RNF212B*, but was not significant in sex-specific regional heritability models, nor the GWAS analyses. Since then, addition of more SNP genotyped individuals and improvements in pedigree construction methodology (Huisman, 2017) now allows us to characterise recombination rate in a further ∼3,900 gametes. Furthermore, advances in genotype imputation methods in complex pedigrees has allowed us to increase the SNP dataset for conducting genome-wide association study from ∼39,000 to ∼417,000 SNPs.

Here, we revisit our previous study to carry out GWAS of male and female recombination rate. Our motivation was to: (a) confirm a sex-limited association with *RNF212* on chromosome 6; (b) fine-map the association on chromosome 7 and determine the action and effect size of this locus in both males and females; and (c) to identify any further associations that could not be detected using the previous dataset.

## Materials and Methods

### Study Population

Soay sheep living within the Village Bay area on the island of Hirta, St Kilda, Scotland (57°49‘N, 8°34‘W) have been individually studied since 1985 (Clutton-Brock *et al*., 2004). All sheep are ear-tagged at their first capture, including 95% of lambs born within the study area, and the majority of animals are recaptured on an annual basis. DNA samples are routinely obtained from ear punches and/or blood sampling. All animal work is carried out according to UK Home Office procedures and was licensed under the UK Animals (Scientific Procedures) Act 1986 (License no. PPL60/4211).

### Genotype and Pedigree Data

All sheep were genotyped at 51,135 SNP markers on the Illumina Ovine SNP50 BeadChip (Kijas *et al*., 2009). Quality control was carried out using the check.marker function in GenABEL v1.8-0 with the following thresholds: SNP minor allele frequency (MAF) > 0.001, SNP locus genotyping success > 0.99, individual sheep genotyping success > 0.95; a total of 7,700 sheep and 39,398 SNPs remained. Pedigree relationships were previously inferred using data from 438 SNP loci in the R package Sequoia v1.02 (Huisman, 2017) and from field observations between mothers and their offspring born within the study area. Autosomal SNP genotypes were used to calculate the 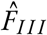 genomic inbreeding coefficients using GCTA v1.90.2 (Yang *et al*., 2011).

A further 189 individuals were genotyped at 430,702 polymorphic SNPs on the Ovine Infinium HD SNP BeadChip for imputation of genotypes into individuals typed on the 50K chip (see Johnston *et al*. 2016 for method and individual selection criteria and Stoffel *et al*. 2020 for full imputation method for this dataset). Briefly, SNP genotypes from the HD Chip were imputed into the SNP50 Chip typed individuals using AlphaImpute v1.98 (Hickey *et al*., 2012; Antolín *et al*., 2017) resulting in a dataset with 7,691 individuals genotyped at 417,373 SNPs, with a mean genotyping rate per individual of 99.5% (range 94.8% - 100%). As there were no X-chromosome SNPs in common between the two SNP Chips, only the SNP50 Chip SNPs were used for association analyses on the X chromosome. SNP positions for both chips were are known relative to sheep genome assembly Oar_v3.1 (GenBank assembly ID GCA_000298735.1).

### Estimation of autosomal crossover counts and linkage maps

Autosomal crossover positions were estimated using an identical protocol to that outlined in Johnston *et al*. (2016). Briefly, the method uses Ovine SNP50 BeadChip SNP data from a focal individual’s parents, mate and offspring to characterise crossovers that occurred in the gametes transmitted from the focal individual to its offspring. Crossovers were determined using the software CRI-MAP v2.504a (Green *et al*., 1990). Here, we estimated ACC for a further 3,908 gametes, leading to a dataset of ACC for 7,235 gametes transmitted from 1,632 unique focal individuals. Because of differences in the SNP quality control between analyses, we used 37,853 SNPs in common with the previous recombination rate study. Simulation studies previously showed that this method will identify >99% of crossovers, meaning that ACC estimated using this method can be used as a proxy for individual recombination rate. The method also provides linkage maps for each chromosome, allowing us to update the existing map for Soay sheep to include information from a much larger number of meioses.

### Animal Models

A restricted maximum likelihood (REML) animal model approach (Henderson, 1975) was used to estimate components of phenotypic variance for ACC, including the additive genetic effect (i.e. heritability). A genomic relatedness matrix (GRM) was constructed for all polymorphic autosomal SNPs from the 50K chip using GCTA v1.90.2 (Yang *et al*., 2011) and was adjusted with the argument *--grm-adj 0*, which assumes a similar frequency spectra of genotyped and causal loci. Related individuals were not removed from the GRM as there is high relatedness in the study population, and there are were no significant parental effects or common environment effects on individual recombination rate (Johnston *et al*., 2016). Animal models were constructed for both sexes combined, females only and males only, and were run in ASReml-R v4 (Butler *et al*., 2009) in R v3.6.2. Fixed effects included sex (in the combined model) and the genomic inbreeding coefficient 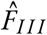 initial models fit age in years fit as a linear covariate, but as it was not significant, it was removed from the final models (Figure 1). The significance of fixed effects was tested using Wald tests. Random effects included the additive genetic effect (as estimated using the GRM) and identity of the focal individual (i.e. “permanent environment”) effect to account for repeated measures. Despite not being significant, the fixed effect 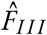 and the random permanent environment effect were retained in all models to account for potential underestimation of ACC due to long runs of homozygosity and pseudoreplication, respectively. Significance of the additive genetic effect was determined by dropping the additive genetic effect from each model and comparing with the full model using a likelihood ratio test distributed as 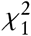. Bivariate models examining the genetic correlation *r*_*A*_ between male and female ACC were run using the CORGH error structure in ASReml-R (correlation with heterogeneous variances). Models were set with *r*_*A*_ to be unconstrained. To test whether *r*_*A*_ was significantly different from 0 and 1, the model was compared to models with *r*_*A*_ fixed at a value of 0 or 0.999 using likelihood ratio tests.

**Figure 1:**
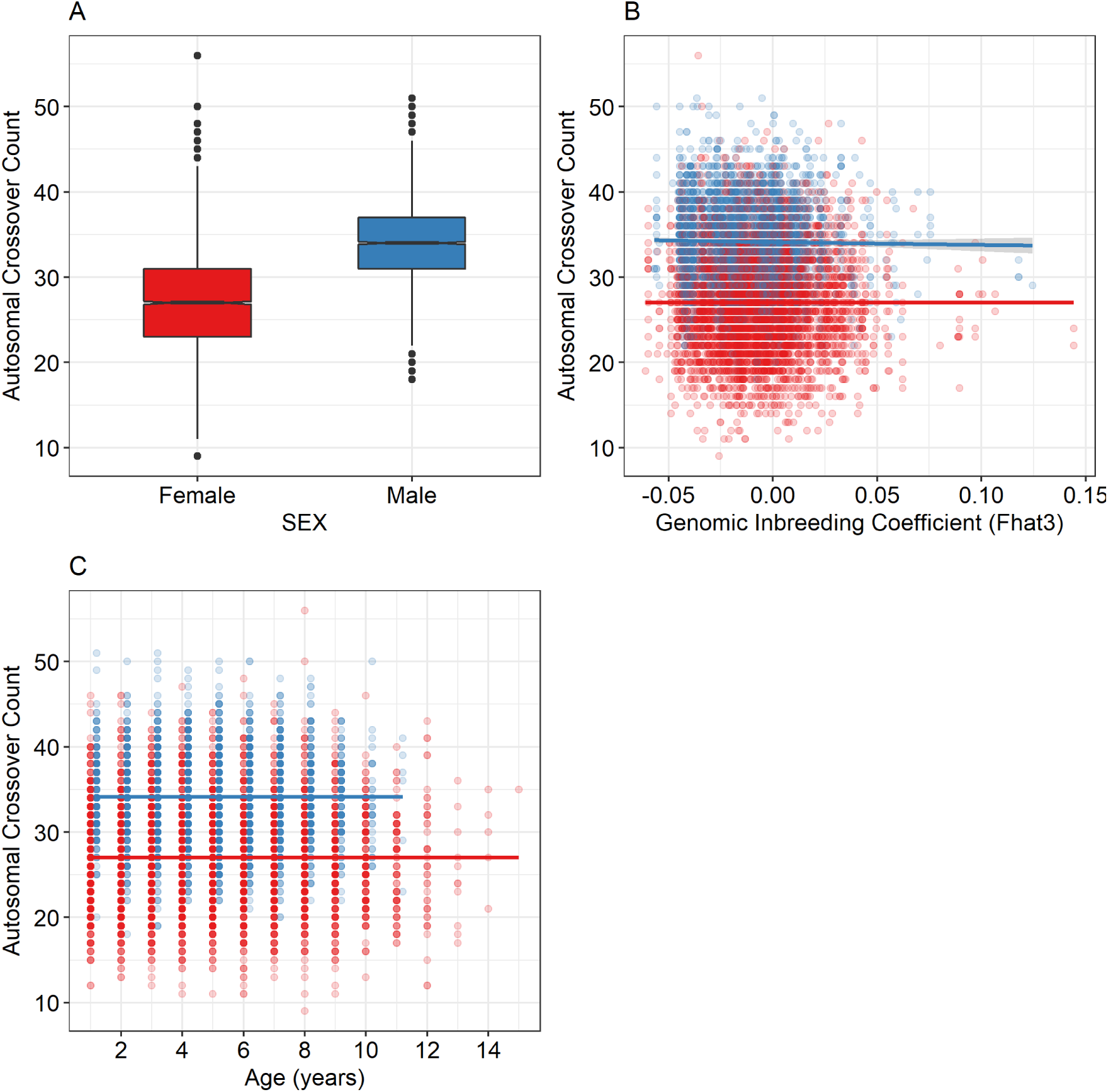
Association between autosomal crossover count and: A. sex; B. inbreeding coefficient 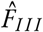; and C. age in years. Plots were made using the raw data. Points are red for females and blue for males. Lines in B. and C. are general additive model smoothing parameters as fit by the plotting library ggplot2 v3.3.2 (Wickham, 2016) in R v3.6.2. Points in plot C are slightly jittered between the sexes on the x-axis to more clearly visualise the sex differences with age. Sample sizes are provided in Table 1.

### Genome-wide association studies

Genome-wide association studies of ACC with the imputed SNP dataset were carried out using function *rGLS* in the package RepeatABEL v1.1.31 (Rönnegård *et al*., 2016) in R v3.6.2. This approach fits both repeated measures and the GRM, the latter accounting for population structure. Association statistics were corrected for any further inflation by dividing by the genomic control parameter *λ*, calculated as the median observed 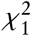 statistic divided by the median 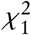 expected under a null distribution. The significance threshold at *α* = 0.05 was calculated by Stoffel *et al*. (2020) as *P* < 1.28 × 10^−06^, using a ‘simpleM’ approach using linkage disequilibrium information to determine the effective number of independent tests (Gao *et al*., 2008).

**Table 1:** Data information and animal model results for autosomal crossover count (ACC) in Soay sheep. Numbers in parentheses are the standard error unless otherwise stated. N_*OBS*_ is the observed number of gametes, with numbers in parentheses the number of unique focal individuals. N_*xovers*_ are the total number of crossovers in the dataset. The mean ACC and the variance (V_*OBS*_) were calculated from the raw data. *V*_*P*_ is phenotypic variance. *h*^2^, *pe*^2^ and *e*^2^ are the narrow-sense heritability, the permanent environment effect, and the residual effect, respectively; all are calculated as the proportion of *V*_*P*_.

| Analysis | $N_{OBS}$ | $N_{xovers}$ | Mean | $V_{OBS}$ | $V_P$ | $h^2$ | $pe^2$ | $e^2$ |
| --- | --- | --- | --- | --- | --- | --- | --- | --- |
| Both Sexes | 7626<br>(1483) | 227646 | 29.851 | 41.32 | 29.514<br>(0.588) | 0.148<br>(0.019) | 0.025<br>(0.013) | 0.827<br>(0.014) |
| Males | 3043<br>(434) | 103914 | 34.149 | 23.974 | 24.162<br>(0.72) | 0.118<br>(0.032) | 0<br>(0.024) | 0.882<br>(0.02) |
| Females | 4583<br>(1049) | 123732 | 26.998 | 32.439 | 32.41<br>(0.797) | 0.181<br>(0.024) | 0<br>(0.017) | 0.819<br>(0.017) |

For the most highly-associated SNPs, genotype effect sizes were estimated by fitting the SNP genotype as a three level factor in the animal models described above. Then, in separate models, the proportion of phenotypic variance explained by significant regions was determined using a regional heritability approach (Nagamine *et al*., 2012). GRMs were constructed as above for the 20 SNPs from the SNP50 chip spanning the highest associated SNP in that region (i.e. 10 SNPs from either side of the association). In one case where the association was at the end of the chromosome, the last 20 SNPs on that chromosome were used to construct the GRM. These regional GRMs were then fit as additional random effects in the animal models described above, with their significance determined using likelihood ratio tests distributed as 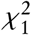.

In order to identify potential candidate genes for ACC, sheep gene IDs SNPs in the broad associated regions were extracted from Ensembl (gene build ID Oar_v3.1.100) using the function *getBM* in the R package biomaRt v2.42.1 (Durinck *et al*., 2009) in R v3.6.2. Gene orthologues in humans (*Homo sapiens*), cattle (*Bos taurus*), mouse (*Mus musculus*) and rat (*Rattus norvegicus*) were extracted using the function *getLDS*. The associated gene ontology (GO annotations) information for all genes and orthologues were then extracted using the *getBM* function. All gene names, phenotype descriptions, GO terms and definitions were then queried for terms associated with meiosis and recombination, using the R command *grep* with the text strings *meio* and *recombin*.

## Data Availability

Raw data will be publicly archived. Code for the analysis is archived at https://github.com/susjoh/2020_Soay_Recomb_GWAS.

## Results

### Variation and heritability of autosomal crossover count

Males had higher recombination rates than females, with 7.24 more autosomal crossovers observed per gamete (SE = 0.18. Wald 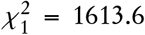, P < 0.001; Figure 1). There was no association with ACC and age or the inbreeding coefficient 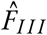 (P > 0.05; Figure 1). Females had higher phenotypic variance than males (female *V*_*P*_ = 32.41, male *V*_*P*_ = 24.162; Table 1). ACC was heritable in both sexes (*h*^2^ = 0.148) and in males and females separately (*h*^2^ = 0.118 and 0.181, respectively; Table 1); likelihood ratio tests confirmed that including the additive genetic effect significantly improved the animal models (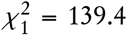, 19.72 and 119.9 for both sexes, males, and females, respectively; P < 0.001). The permanent environment contribution to phenotypic variance was not significant in any models (P > 0.05), with the remaining variance attributed to residual effects (Table 1). Bivariate models of ACC between males and females showed a positive genetic correlation (*r*_*A*_ = 0.555, SE = 0.138) that was significantly different from both *r*_*A*_ = 0 and 1 (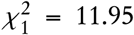 and 17.91, respectively; P < 0.001). Linkage map lengths from the ACC analysis in this study were strongly correlated with the previous maps (linear regression, adjusted *R*^2^ >0.999, Figure S1). The updated map had 15,787 unique centiMorgan positions compared with 12,359 in the previous study; it is provided in Table S2 and Figure S2.

### Genome-wide Association Study

There was a strong association between SNPs at the sub-telomeric region of chromosome 6 and female ACC (*oar3_OAR6_116402578*, P = 2.45 × 10^−17^; Figures 2 & 4; Tables 2 & S3). This locus explained 45.9% of the heritable variation in female ACC, with a difference of 5 crossovers per gamete between the two homozygotes (Figure 3, Table 2). There was no significant association with this locus and male ACC, confirming that its action is likely to be sex-limited. The same SNP showed the highest genome-wide association in the previous study (Johnston *et al*., 2016) and occurs 25.2 kb from the putative location of *RNF212*. It should be noted that *RNF212* is not annotated on the Oar_v3.1 sheep genome due to a gap in the sequence, but is predicted to be positioned between 116,427,802 and 116,446,904 bp (Johnston *et al*., 2016); *RNF212* is present in this position on the Oar_rambouillet_v1.0 genome assembly (GenBank assembly ID GCA_002742125.1). A second association was observed in both sexes on chromosome 7 (*oar3_OAR7_21368818* and *oar3_OAR7_21347355*, P = 6.11 × 10^−13^, 1.83 × 10^−10^ and 7.20 × 10^−7^ in both sexes combined, females and males, respectively); the association statistic in males was also identical for locus *oar3_OAR7_21116299* (Figures 2 & 5; Tables 2 & S3). The action of this locus was similar between males and females, with a difference of ∼4.5 crossovers per gamete between the two homozygotes and explaining between 19.7% and 24.8% of the heritable variation in ACC (Figure 3, Table 2). These SNPs spanned the candidate locus *RNF212B* (21,241,831 to 21,273,165 bp). GO term analysis indicated that both *RNF212* and *RNF212B* were the closest genes to each hit that were associated with meiotic processes (Table S4; Figures 4 & 5).

**Table 2:**
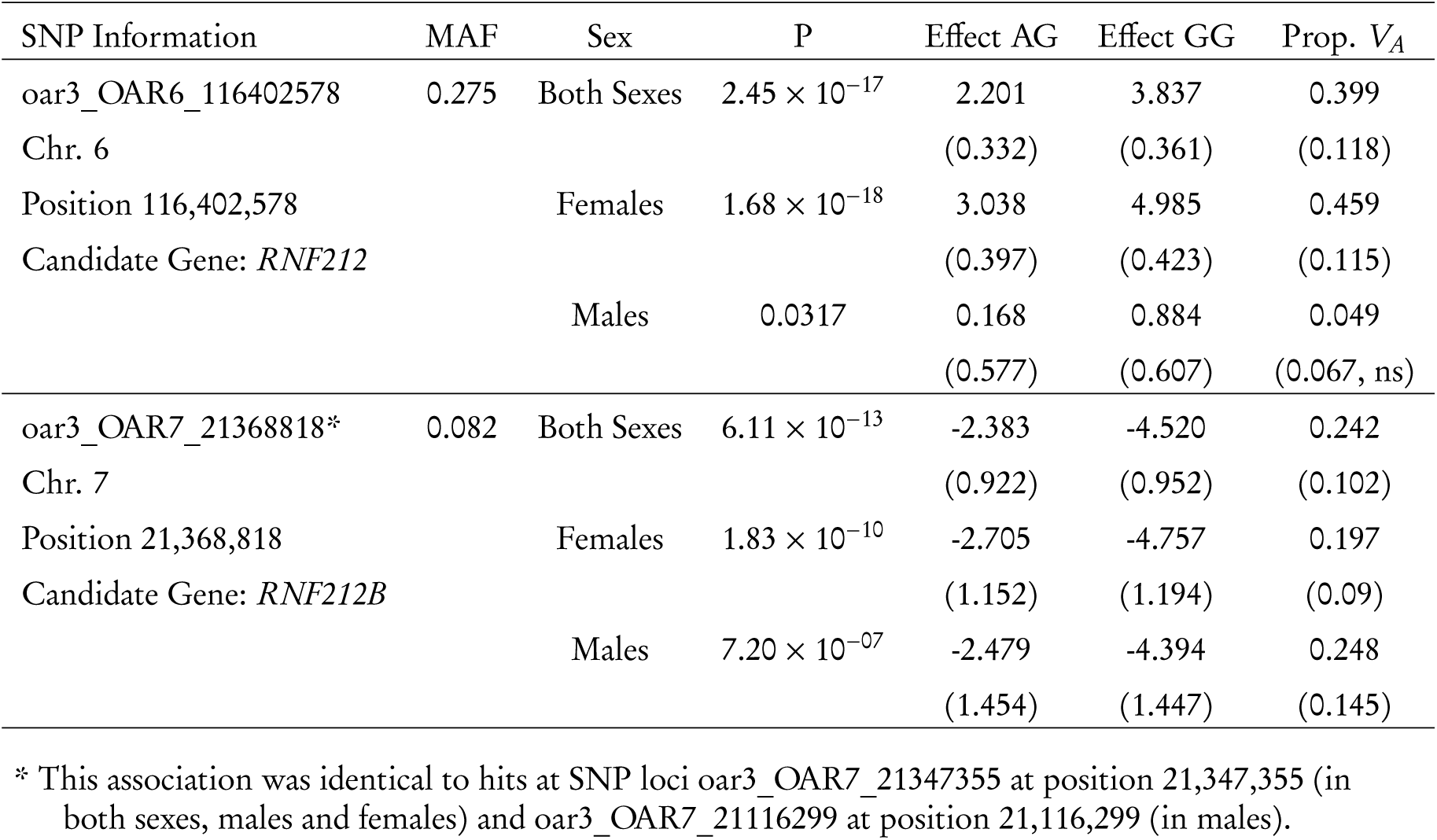
The most highly associated SNPs from GWAS of ACC in Soay sheep. MAF is the minor allele frequency (corresponding to the A allele for each locus). Sex indicates if the model was run in both sexes combined, or in females or males only. P is the P-value for the association statistic after correction using genomic control *λ*. Effect sizes are provided for animal models where the genotype fit as a fixed factor, and are given relative to the model intercept of 0 at genotype AA. Prop. *V*_*A*_ is the proportion of the additive genetic variance explained by the genomic region containing the SNP locus. All values in parentheses are standard errors. Full GWAS results are provided in Table S3. Effect sizes and associated sample sizes for each genotype are shown in Figure 3.

**Figure 2:**
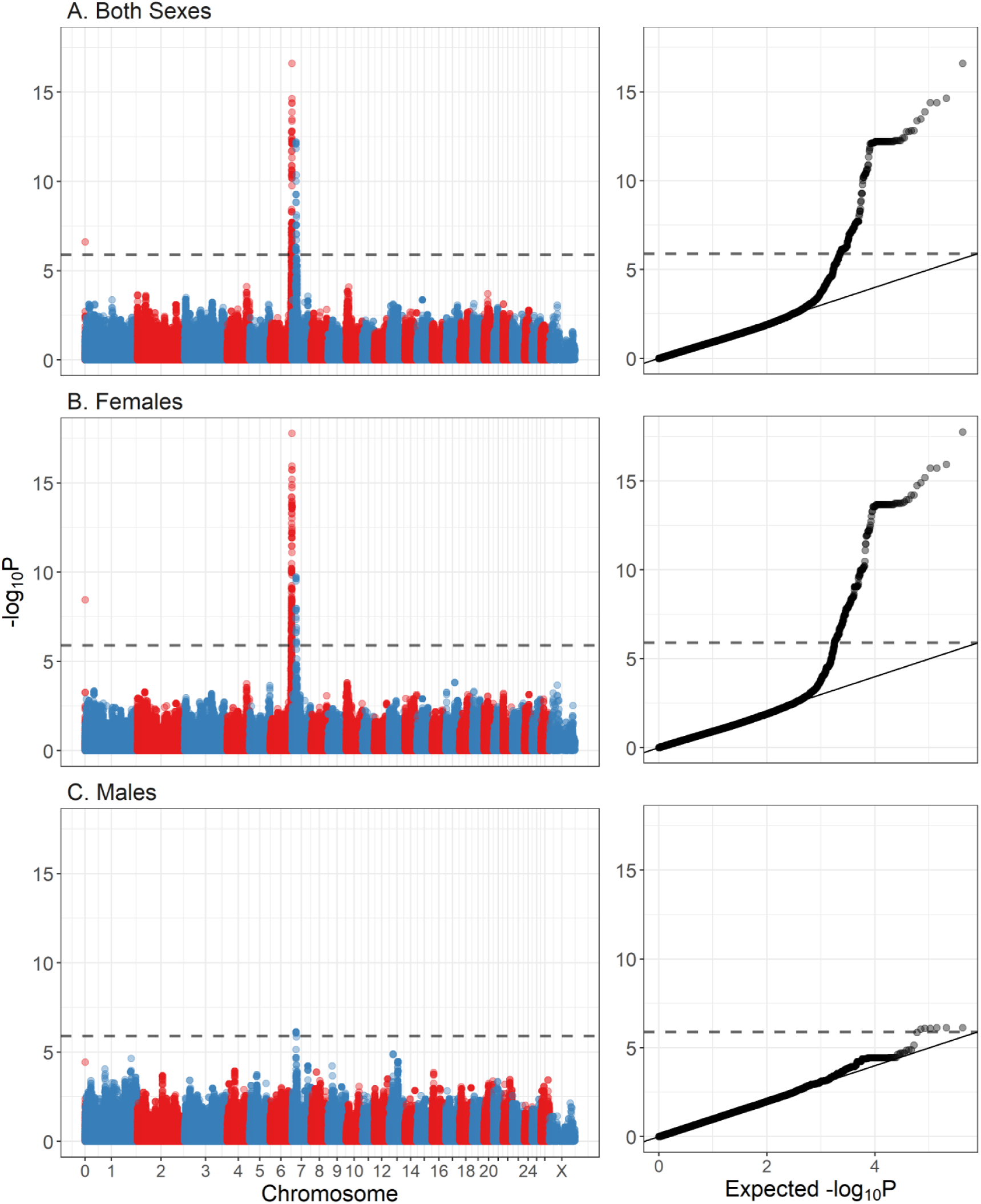
Genome-wide association plots of autosomal crossover counts for A. both sexes combined, B. females only and C. males only. The dashed line is the significant threshold equivalent to *α* <0.05. The left panels show the association for individual SNPs relative to their genomic position, with points colour-coded by chromosome. The right plots show the distribution of the observed *log*_10_*P* values against those expected under a null distribution. All association statistics have been corrected for genomic control using the *λ* parameter. All results are provided in Table S3. Sample sizes are provided in Table 1.

**Figure 3:**
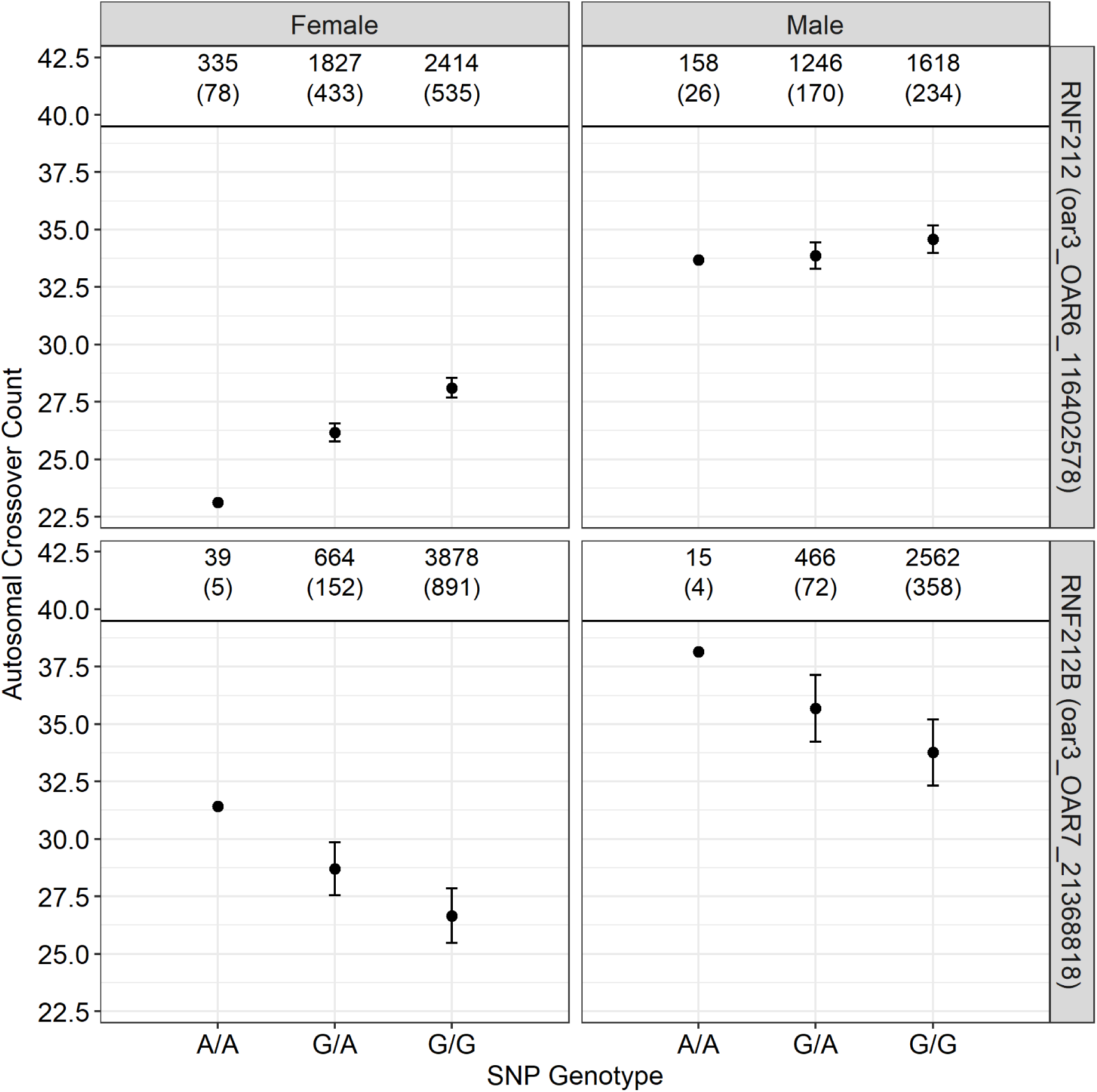
Effect sizes for the most highly associated SNPs shown for males and females, as calculated from animal models including the SNP genotype as a fixed factor. Genotype AA is the model intercept. Numbers indicate the number of observations at each genotype, with the number of unique focal individuals in parentheses.

**Figure 4:**
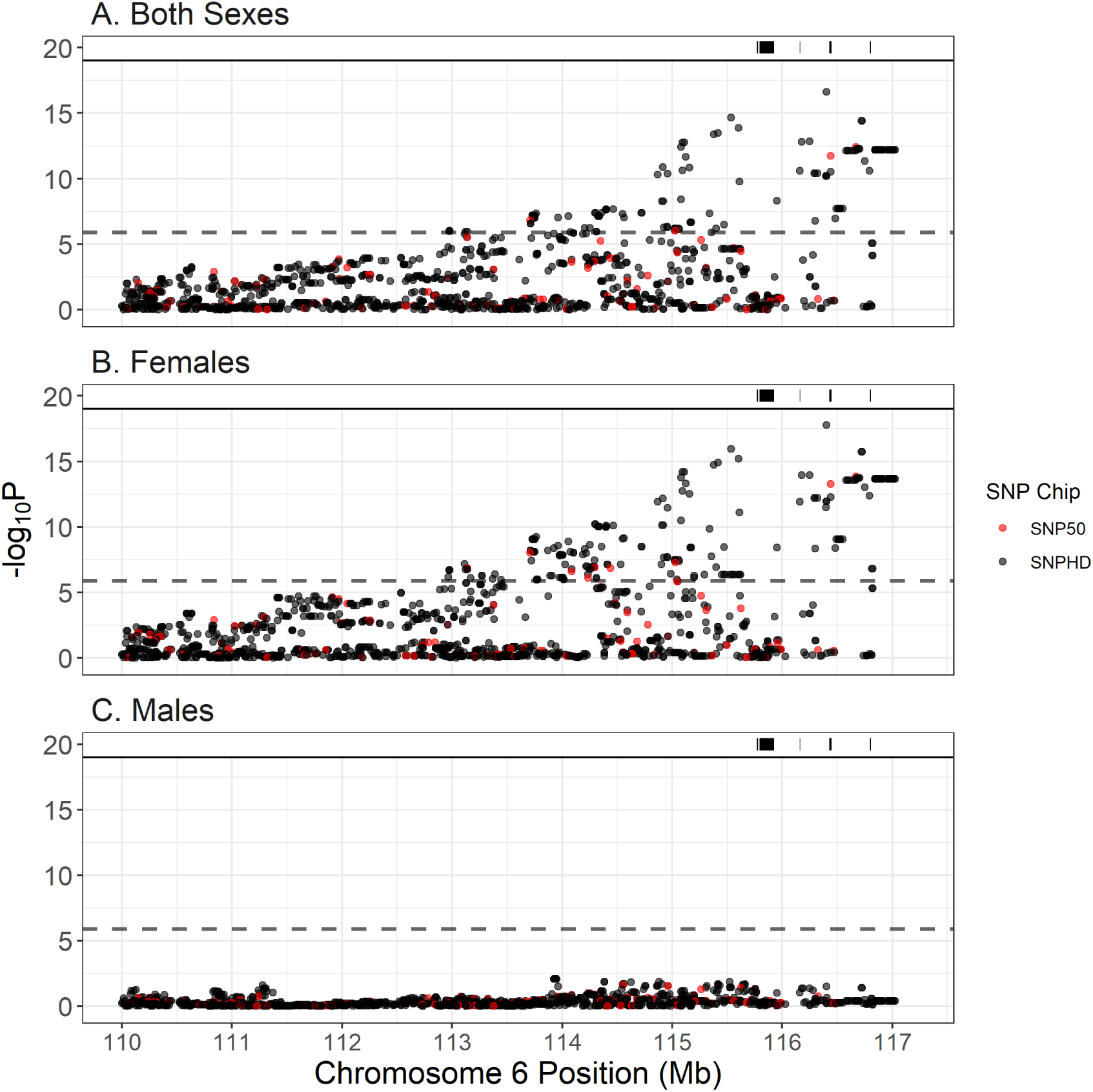
Regional association statistics for the significant region on chromosome 6. Each point represents a single SNP locus, coloured by their origin on the SNP50 (red) or SNPHD Chips (black, i.e. imputed genotypes). The positions of genes associated with meiotic processes are shown as black bars at the top of each plot (Gene information in Table S4).

## Discussion

This study revisited a previous analysis investigating the genetic architecture of ACC in Soay sheep, with more than double the number of gametes and more than ten times the number of SNP loci. We confirmed that ACC is ∼1.27 times higher in males than in females, and is not associated with age or inbreeding in neither males nor females. We identified two candidate genes associated with ACC: *RNF212* on chromosome 6, which is associated with female recombination rate only; and its paralogue *RNF212B* on chromosome 7, which is associated with recombination rate in both sexes. Both loci have repeatedly been associated with recombination rate variation in mammals, with all studies conducted to date implicating either one or both of these loci; these include humans (Kong *et al*., 2008, 2014), cattle (Sandor *et al*., 2012; Ma *et al*., 2015; Kadri *et al*., 2016), pigs (Johnsson *et al*., 2020), deer (Johnston *et al*., 2018) and domestic sheep (Petit *et al*., 2017). Functional studies in mice indicate that the protein RNF212 is essential for crossover formation during meiosis, and that it has a dosage sensitive effect (Reynolds *et al*., 2013). We observe an additive effect of alleles on ACC at both loci in this study, suggesting that a similar dosage sensitive effect is the mechanism driving rate differences in Soay sheep.

The current study confirms the previous association at *RNF212* and does not gain any novel insights into the nature of its association with ACC. However, the observation at *RNF212B* builds on previously tentative evidence of an association in this genomic region. In the 2016 study, a genome-wide regional heritability analysis identified an association in both sexes combined in a 1.09Mb region, which contained *RNF212B* and another candidate locus, meiotic recombination protein *REC8*. However, there was no corresponding association using GWAS and there was no significant effect observed in the sex-specific regional heritability analyses, providing limited insight into the action of this region on ACC variation. Here, we have shown that adding a larger number of gametes and SNPs has increased the power to detect an association at *RNF212B*, and indicating that ACC variation is more likely to be attributed to *RNF212B* than *REC8* (although see further discussion below). The MAF of *RNF212B* is relatively low (0.082), suggesting that lower sample sizes in the 2016 study were not sufficient to detect the effects of rare loci within the population. Using the SNP50 Chip alone, the increased number of gametes in current study would have shown a significant hit on chromosome 7 in females (but not in males; Figure 5), however, the highest association would be around 345Kb away from the that observed in the current study. Our work shows that increasing marker density using imputation is likely to be an increasingly important approach in conducting GWAS in natural populations.

**Figure 5:**
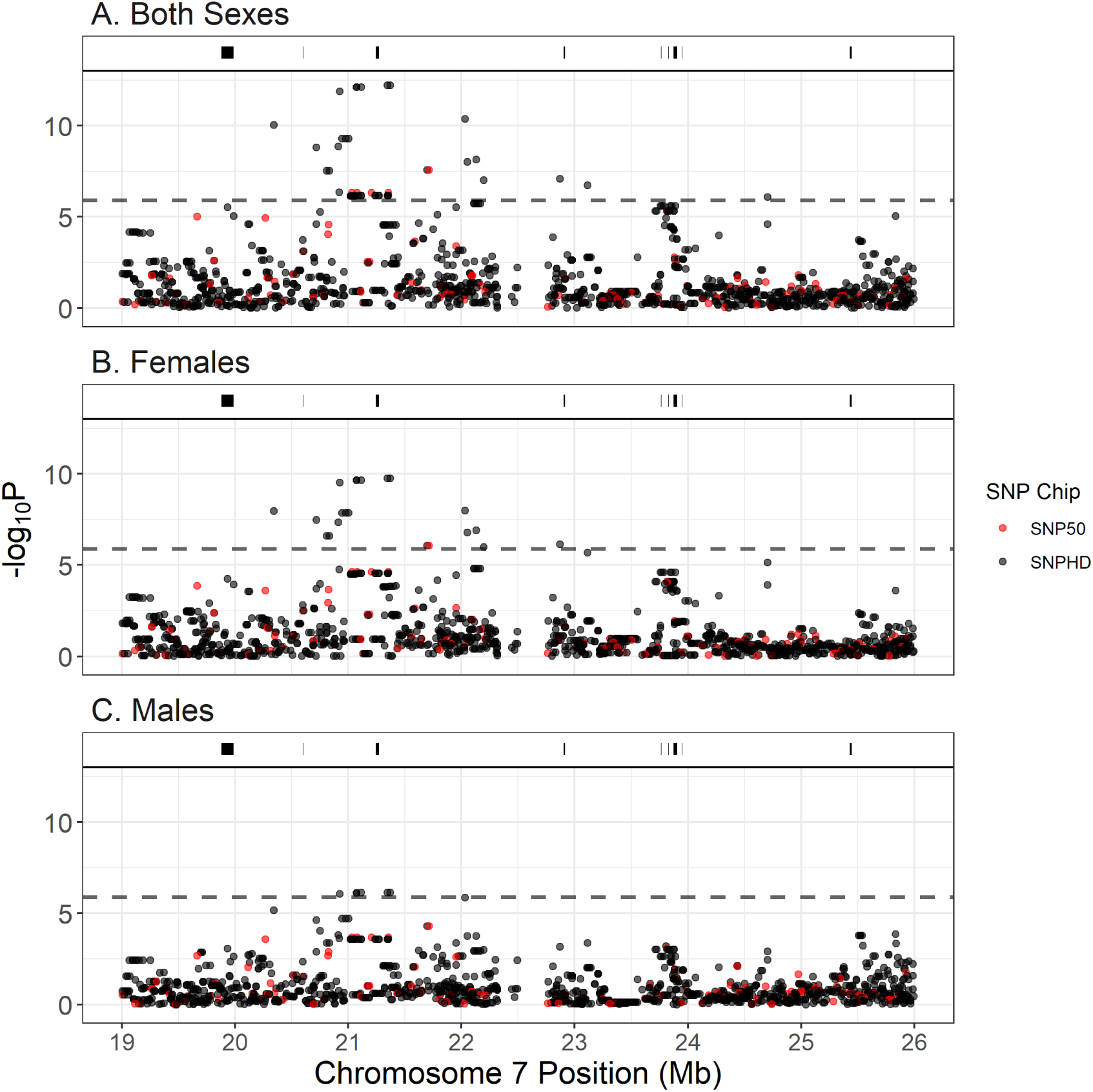
Regional association statistics for the significant region on chromosome 7. Each point represents a single SNP locus, coloured by their origin on the SNP50 (red) or SNPHD Chips (black, i.e. imputed genotypes).The positions of genes associated with meiotic processes are shown as black bars at the top of each plot (Gene information in Table S4).

Soay sheep have extensive linkage disequilibrium (LD) throughout the genome due to recent population bottlenecks, a small effective population size and a high prevalence of inbreeding (Clutton-Brock *et al*., 2004; Bérénos *et al*., 2014; Stoffel *et al*., 2020). An advantage of this is that there is a higher chance of typing SNP loci that are in LD with causal loci, and that genotype imputation can be carried out with high accuracy, increasing the likelihood of detecting associated regions. However, a disadvantage is that high LD within the population makes it difficult to separate the effects of linked loci that may also contribute to phenotypic variation in the population. The associated regions on both chromosomes 6 and 7 extend over several megabases, with several other loci associated with meiotic processes occurring within these regions (Figures 4 & 5, Table S4); we cannot rule out that these loci also affect variation in ACC in Soay sheep. Functional validation of the role of *RNF212* and *RNF212B* is not possible within this wild system as experimental manipulation and invasive sampling are prohibited. However, further studies in domestic sheep (e.g. Petit *et al*. 2017) and related species may further elucidate the role of loci within these regions and the relative importance of regulatory and protein-coding variation in driving recombination rate variation.

Overall, this study has highlighted the merits of genotype imputation and increasing sample sizes to determine the genetic architecture of recombination rate variation in a natural system. Identifying candidate loci and their sex-specific effect sizes provides a stepping stone for future studies investigating the evolution of recombination rates within this and related systems, through modelling the temporal dynamics of these loci, their origin and their association with individual fitness.

## Supporting information

Supplemental Table S2 - Soay sheep Linkage Map.

Supplemental Table S3 - GWAS Results for Gamete Autosomal Crossover Count

Supplemental Table S4 - Genes in Regions Associated with Gamete Autosomal Crossover Count

A file containing Supplemental Figures S1 & S2, Table S1 and descriptions of Tables S2-S4.

## Acknowledgements

We thank Jill Pilkington, Ian Stevenson and all Soay sheep project members and volunteers for collection of data and samples. Camillo Bérénos, Philip Ellis, Hannah Lemon and Penny Jack prepared DNA samples for Ovine SNP50 BeadChip genotyping and Jisca Huisman assisted with the pedigree construction. Genotyping was performed by Lee Murphy and colleagues at the Wellcome Trust Clinical Research Facility Genetics Core in Edinburgh. This work has made use of the resources provided by the Edinburgh Compute and Data Facility (http://www.ecdf.ed.ac.uk/). Permission to work on St Kilda is granted by The National Trust for Scotland, and logistical support is provided by QinetiQ and Eurest. The Soay sheep project has been supported by grants from the UK Natural Environment Research Council, with the SNP genotyping funded by a European Research Council Advanced Grant to JMP. SEJ is supported by a Royal Society University Research Fellowship.

## Author Contributions

S.E.J and J.M.P. conceived the study. J.M.P organised the collection of samples. M.S. conducted the genotype imputation. S.E.J. analysed the data and wrote the paper. All authors contributed to revisions.

## Notes

### Competing Interest Statement

The authors have declared no competing interest.

