## Supplementary material for "Variants at *RNF212* and *RNF212B* are associated with recombination rate variation in Soay sheep (*Ovis aries*)": A file containing Supplemental Figures S1 & S2, Table S1 and descriptions of Tables S2-S4.

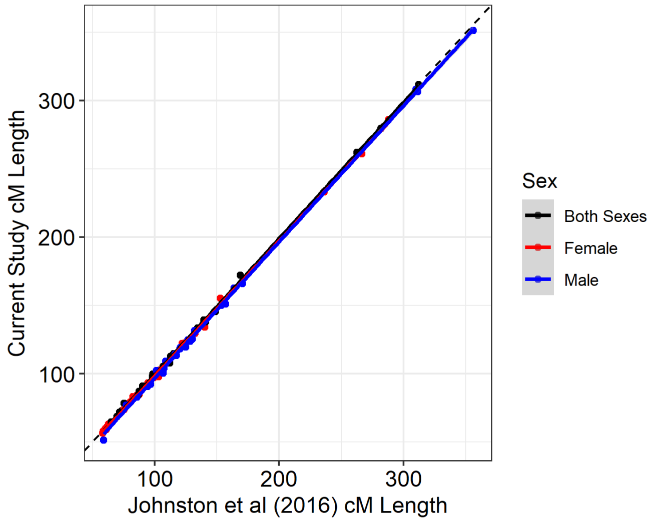


**Figure S1:** Correlation between linkage map lengths determined in the previous study (Johnston *et al.*, 2016) and in the current study. Each point indicates a chromosome, and lines indicate linear regressions. Points are coloured by sex-averaged or sex-specific map types. The black dashed line indicates a perfect correlation with slope = 1 and intercept = 0.

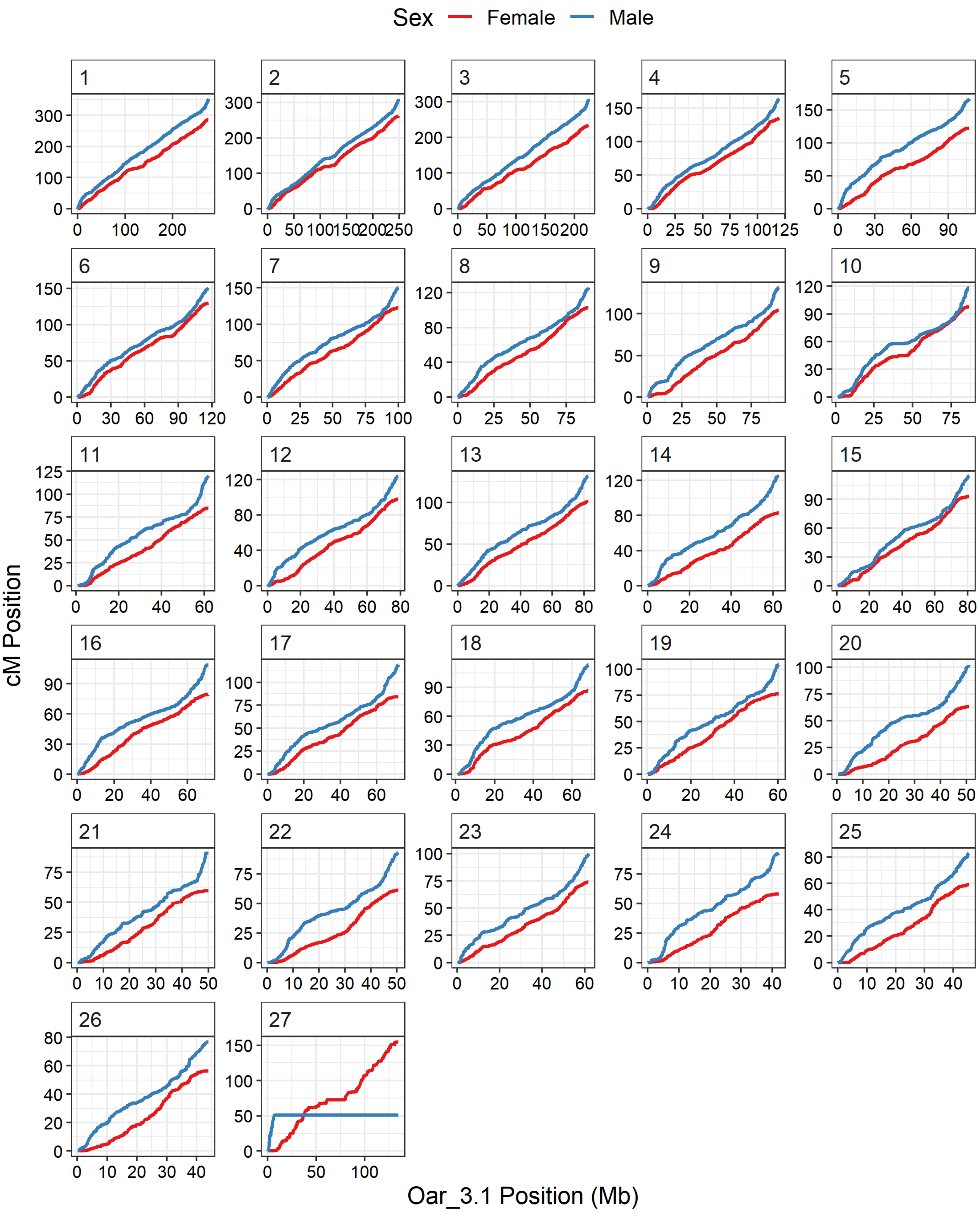


**Figure S2:** Sex-specific linkage maps for each sheep chromosome plotted relative to the sheep genome assembly Oar_v3.1. The underlying map data is provided in Table S2. Chromosome 27 is the X chromosome, with the short map segment in male sheep corresponding to the pseudoautosomal region (PAR).

**Table S1:** Fixed effects in animal models of autosomal crossover count. Models were run in both sexes, males and females. All Wald statistics were equivalent to a $\chi^2$ test with 1 degree of freedom.

| Model | Effect | Estimate | SE | Z Ratio | Wald Statistic | P |
|---|---|---|---|---|---|---|
| Both Sexes | (Intercept) | 26.966 | 0.106 | 253.849 | 112596.078 | 0.0000 |
| | Fhat3 | 1.148 | 3.692 | 0.311 | 0.097 | 0.7557 |
| | Sex (Male) | 7.236 | 0.180 | 40.161 | 1613.634 | 0.0000 |
| Males | (Intercept) | 34.178 | 0.144 | 236.710 | 61239.042 | 0.0000 |
| | Fhat3 | -3.402 | 5.872 | -0.579 | 0.336 | 0.5623 |
| Females | (Intercept) | 26.970 | 0.113 | 238.303 | 59962.846 | 0.0000 |
| | Fhat3 | 0.853 | 4.610 | 0.185 | 0.034 | 0.8533 |

**Table S2:** Linkage map information for the current study. Order is the order of markers on the chromosome. SNP.Name is the Ovine SNP50 BeadChip identifier. Chr is the chromosome number, where 27 is the X chromosome. cMPosition, r, cMdiff are the sex-averaged centiMorgan position, recombination fraction and centiMorgan difference with the following locus, respectively; similarly this is provided for Female and Male maps. Oar3_Chr and Oar3_Pos are the relative chromosome number and position relative to the sheep genome (Oar_v3.1), cMPosition.2016, cMPosition.Female.2016 and cMPosition.Male.2016 indicate the centiMorgan positions as determined by the previous study (Johnston *et al.*, 2016).

[File: Table_S2_Linkage_Map.txt]

**Table S3:** Association statistics for genome-wide association studies of autosomal chromosome counts. Sex indicates analyses for both sexes, females only and males only. SNP.Name is the Ovine SNP50 BeadChip identifier. Chromosome and Position (bp) are given relative to the sheep genome Oar_v3.1. A1 and A2 reference and alternate allele at each SNP. effB is the slope of the effect of allele A2, with the standard error se_effB. Chi2.1df and P1df is the association chi-squared statistic and associated P-value, respectively, before correction with genomic control $\lambda$. Pc1df is the corrected P-value after genomic control. Exp is the corresponding P-value for that SNP locus assuming a null distribution of P-values (see Figure 2). Q.2 is the minor allele frequency. Cumu is the cumulative genomic distance.

[File: Table_S3_ACC_GWAS.txt]

**Table S4:** Gene positions in the GWAS significant regions obtained using biomaRt v2.42.1. ensembl_gene_id is the Ensembl identifier for the sheep gene. external_gene_name is the sheep gene name. chromosome_name, start_position and end_position are the chromosome number, start and stop positions for each gene, respectively. Meiotic indicates whether this gene and/or its orthologues in other species have GO terms associated with meiotic processes. Orthos indicates the unique gene names associated with orthologues of each gene.

[File: Table_S4_Gene_List_Table.txt]
